# Leveraging Family History in Case-Control Analyses of Rare Variation

**DOI:** 10.1101/665075

**Authors:** Claudia R. Solis-Lemus, S. Taylor Fischer, Andrei Todor, Cuining Liu, Elizabeth J. Leslie, David J. Cutler, Debashis Ghosh, Michael P. Epstein

## Abstract

Standard methods for case-control association studies of rare variation often treat disease outcome as a dichotomous phenotype. However, both theoretical and experimental studies have demonstrated that subjects with a family history of disease can be enriched for risk variation relative to subjects without such history. Assuming family history information is available, this observation motivates the idea of replacing the standard dichotomous outcome variable used in case-control studies with a more informative ordinal outcome variable that distinguishes controls (0), sporadic cases (1), and cases with a family history (2), with the expectation that we should observe increasing number of risk variants with increasing category of the ordinal variable. To leverage this expectation, we propose a novel rare-variant association test that incorporates family history information based on our previous GAMuT framework (Broadaway et al., 2016) for rare-variant association testing of multivariate phenotypes. We use simulated data to show that, when family history information is available, our new method outperforms standard rare-variant association methods like burden and SKAT tests that ignore family history. We further illustrate our method using a rare-variant study of cleft lip and palate.

## 1 Introduction

Sequencing and exome-chip technologies facilitate the discovery of rare genetic variation influencing complex diseases. Many rare-variant association studies of complex diseases now exist with most studies employing traditional case-control sampling designs for analysis (De Rubeis et al., 2014; Sanders et al., 2017). Under such a design, studies typically test whether patterns of rare variation within a gene or region of interest differ between affected and unaffected subjects using either burden (Li and Leal, 2008) or variance-component (Wu et al., 2011) approaches based on an underlying logistic-regression framework that treats disease status as a simple dichotomous outcome variable. While such an analysis strategy is commonplace, there may exist helpful secondary information collected by the study that can facilitate the creation of a modified outcome variable that is more refined than the coarse dichotomous outcome typically considered. Use of this refined outcome variable within the study can reduce heterogeneity and potentially lead to more powerful analyses.

One valuable source of secondary information often collected in a case-control study (but rarely utilized) is whether a sample participant reports a family history of the disease under study. Subjects with a family history of disease demonstrate different patterns of genetic variation than their sporadic counterparts. In particular, several papers have noted that a sample of cases reporting affected relatives are more enriched for a causal variant than cases without such family history (Teng and Risch, 1999; Zöllner, 2012; Epstein et al., 2015) since more risk variants tend to segregrate in families with multiple affected individuals. Likewise, controls with a family history of disease should have elevated frequency of a causal variant compared to sporadic controls (Liu et al., 2017). These observations motivate replacement of the standard dichotomous outcome variable for disease with a more refined variable that incorporates family-history information into the coding.

In deciding how to refine the variable, we note that we should expect the frequency of a risk variant to follow a gradient that increases in frequency from sporadic controls to controls with a family history to sporadic cases to cases with a family history. One way to exploit this phenomenon in genetic analysis is to recode the disease variable as a ordinal cateogorical variable with four possible levels: controls (0), controls with a family history (1), sporadic cases (2), and cases with a family history (3). If family-history information is unavailable for controls, we instead consider a ordinal categorial variable with three possible levels: controls (0), sporadic cases (1), and cases with a family history (2). In either case, this recoding requires the development of novel methods for rare-variant analysis that can handle ordinal variables. To fill this gap, we propose a novel approach that is an extension of our previous GAMuT approach (Broadaway et al., 2016), which is a nonparametric association test using a kernel-distance covariance (KDC) framework that can handle multi-dimensional genotypes and phenotypes. Kernel-based approaches have found success in rare variant associations due to the natural incorporation of epistatic effects, and sparsity in the methodology. Here, we show how GAMuT can model ordinal outcomes in rare-variant analysis while correcting for confounding covariates such as population stratification. Furthermore, just like the standard GAMuT, the newly proposed ordinal GAMuT produces analytical p-values, which facilitates scaling to genome-wide analyses.

The structure of this paper is as follows: after introducing the ordinal GAMuT method using the KDC framework (Gretton et al., 2008; Székely et al., 2007; Kosorok, 2009; Zhang et al., 2012; Hua and Ghosh, 2015), we present simulation work to show that leveraging family history information via ordinal categorical variables can improve power in rare-variant association tests compared to standard dichotomous modeling of disease phenotypes that ignore such information, like the burden test (Li and Leal, 2008) and Sequence Kernel Association Test (SKAT) (Wu et al., 2011). Finally, we apply ordinal GAMuT to rare and less-common variant data from a genome-wide study of craniofacial defects (Leslie et al., 2016a,b; Mostowska et al., 2018).

## 2 Materials and Methods

### 2.1 Leveraging Family Information through Ordinal Phenotype

We assume a sample of *N* subjects that are genotyped for *V* rare variants in a target gene or region, so that *G*_*j*_ = (*G*_*j,1*_, *G*_*j,2*_, …, *G*_*j,V*_) represents the genotypes of subject *j* at *V* rare-variant sites in the gene of interest. Note that *G*_*j,v*_ represents the number of copies of the minor allele that the subject possesses at the *v*^*th*^ variant. Thus, the matrix of rare-variant genotypes for the sample is denoted **G** ∈ ℝ^*N* ×*V*^.

Let **Q** be an *N* -dimensional vector with binary disease status for *N* subjects. That is, *Q*_*j*_ = 0 if subject *j* is a control, and *Q*_*j*_ = 1 if subject *j* is a case. When family history information is available, we can instead employ a more informative ordinal phenotype. Assuming family history information is only available on cases, we can define the ordinal score as 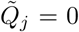 if the subject is a control, 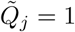 if the subject is a case without family history of the disease, and 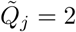 if the subject is a case with family history of the disease. If family history information is available for controls, we can modify appropriately by extending the ordinal variable to the case of four categories: controls without 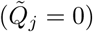 and with family history 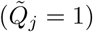, and cases without 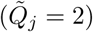 and with family history 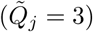. The resulting phenotype vector 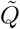 is an *N* -dimensional ordinal vector with disease binary status adjusted for family history for the *N* subjects.

### 2.2 Adjusting for Covariates

After transforming the binary phenotype to ordinal phenotype by incorporating the family history information, we can account for other covariates by regressing the phenotypes 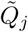 on covariates *X*_*j*_ with a cumulative-logit regression model, and use the residuals in our subsequent rare-variant association test. To illustrate the cumulative-logit regression model, let 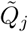 be an ordinal response with *M* categories, and let 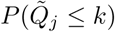 be the cumulative probabilities for *k* = 1, …, *M*. The proportional odds model (McCullagh and Nelder, 1989) is a subclass of cumulative-logit regression models and it is defined as

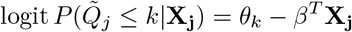

for *k* = 1, …, *M* − 1. Note that the negative sign is a convention to guarantee that large values of *β*^*T*^ **X**_**j**_ increase the probability in the larger values of *k*. In addition, the vector of intercepts *θ* = (*θ*_1_, …, *θ*_*M*− 1_) should satisfy *θ*_1_ *≤ θ*_2_ ≤ … *θ*_*M*−1_.

This model is denoted proportional odds because the ratio of the odds of 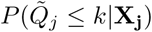 and 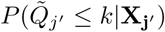 do not depend on the specific category *k*. That is,

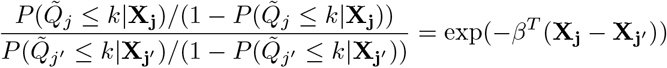

This is also denoted a *parallelism assumption* on *β* (Yee, 2010).

Note that for an ordinal response with *M* categories, we fit *M −* 1 logit regression models. Thus, in our particular setting, we have three categories: controls (*k* = 0), cases without family history (*k* = 1) and cases with family history (*k* = 2), and thus, we will fit 2 models: 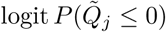 and 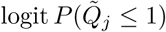. With these models, we estimate the multinomial response probabilities for each individual. That is, for individual *j*, we have:

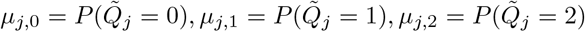

Thus, the matrix of fitted values (denoted **M**) will be a *N* × 3 matrix where each row sums to 1, and the *i*^*th*^ row corresponds to the estimated multinomial probabilities for individual 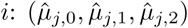. To obtain the matrix of residuals, we first transform the ordinal response into a *N* × 3 binary matrix (denoted 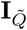) where the *i*^*th*^ row corresponds to the 3-dimensional vector for individual *i* with three indicator functions, one for each category: 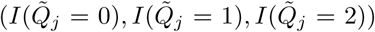. For example, if 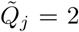, then the binary vector in the *j*^*th*^ row would be (0,0,1). As a result, the matrix of residuals **R** will be the *N* × 3 matrix of the difference between the binary matrix and the matrix of estimated multinomial probabilities: 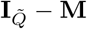. This matrix of residuals will then be input into the GAMuT framework to enable rare-variant association testing. The GAMuT framework allows for correlated phenotypes, and will be described in the following section.

### 2.3 GAMuT Test of Cross-Phenotype Associations

GAMuT tests for independence between the phenotype matrix 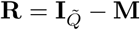 (the *N* × 3 matrix of phenotype residuals) and **G** (the *N* × *V* matrix of multivariate rare-variant genotypes) by constructing an *N* × *N* phenotypic-similarity matrix **Y**, and an *N* × *N* genotypic-similarity matrix **X**. These similarity matrices depend on a user-selected kernel function (Kwee et al., 2008; Schaid, 2010; Wu et al., 2010, 2011). For example, the matrix **Y** can be modeled with the projection matrix: **Y** = **R**(**R**^*T*^**R**)^−1^**R**^*T*^. Alternatively, if *γ*(**R**_*i*_, **R**_*j*_) denotes the kernel function between subjects *i* and *j*, the linear kernel is defined as 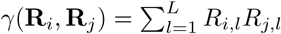 which corresponds to the (*i, j*) entry in **Y**: *Y*_*ij*_. See Broadaway et al. (2016) for more details on other kernel functions to model pairwise similarity or dissimilarity.

After constructing the similarity matrices **Y** and **X**, we center them as **Y**_*c*_ = **HYH** and **X**_*c*_ = **HXH**, where **H** = (**I** − **11**^*T*^ /*N*) is a centering matrix (**HH** = **H**), **I** ∈ ℝ^*N* ×*N*^ is an identity matrix, and **1** ∈ ℝ^*N* × 1^ is a vector of ones. With the centered similarity matrices (**Y**_*c*_, **X**_*c*_), we construct the GAMuT test statistic as

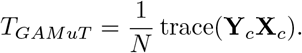

Under the null hypothesis where the two matrices are independent, *T*_*GAMuT*_ follows the asymptotic distribution as the weighted sum of independent and identically distributed 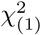 variables (Broadaway et al., 2016). We then use Davies’ method (Davies, 1980) to analytically calculate the p-value of *T*_*GAMuT*_.

### 2.4 Simulations

We conducted simulations to show that ordinal GAMuT properly preserves the type I error and to assess the power of ordinal GAMuT relative to standard case-control burden (Li and Leal, 2008) and SKAT (Wu et al., 2011) tests that do not account for family history information.

For the genetic data, we simulated trios (parents and offspring) with 10,000 haplotypes of 10 kb in size using COSI (Schaffner et al., 2005), a coalescent model that accounts for linkage disequilibrium (LD) pattern, local recombination rate, and population history for individuals of European descent. We defined rare variants as those with *M AF* ≤ 3%. For the power simulations, we assumed the proportion of causal variants to be 15%, with effect size for each causal variant given by 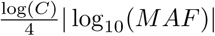 plus a Normal noise with mean 0 and variance 0.1. We varied *C* = 4,6. This setup defines the effect size of any given causal variant as inversely proportional to its MAF, which implies that very rare variants will have a larger effect size.

For the ordinal phenotype, the proband’s probability of disease depended on the sequence data and the disease prevalence, which we varied as 0.01, or 0.05, while the family members’ probability of disease depended on the sequence data and the conditional recurrence risk ratio (*λ* = 2,4,8) (Epstein et al., 2015). If the proband was unaffected, we defined the person as a control. If the proband was affected and none of the parents were affected, we defined the person as a case without family history. Finally, if the proband was affected and at least of the parents was affected, we defined the person as a case with family history.

For each simulated dataset, we generated an equal number of controls, cases without family history, and cases with family history. We varied this number among *N* = 400,750,1000,1500. For each simulated dataset, we applied our ordinal GAMuT method that modeled cases with and without family history separately. We also applied standard burden and SKAT tests that combined all cases together without regards to family history information. For each method, we weighted rare variants using the weighting scheme recommended by Wu et al. (2011); *w*_*v*_ = *Beta*(*M AF*_*v*_, 1,25)*/Beta*(0,1,25).

We used the R package `VGAM` and function `vglm` to fit the cumulative-logit regression model with proportional-odds assumption (Yee, 2010), and use the resulting residuals to construct the phenotypic similarity matrix input in the GAMuT package (Broadaway et al., 2016).

### 2.5 Analysis of Pittsburgh Orofacial Cleft Multiethnic GWAS

Orofacial clefts (OFCs) such as cleft lip (CL), cleft palate (CP), and cleft lip with cleft palate (CLP) are among the most common birth defects in humans with prevalence between 1 in 500 and 1 in 2,500 live births (Tessier, 1976; Mossey et al., 2009). Extensive recent studies identified common nucleotide variants associated with orofacial clefts, such as 1p22.1, 2p24.2, 3q29, 8q24.21, 10q25.3, 12q12, 16p13.3, 17q22, 17q23, 19q13, and 20q12 (Birnbaum et al., 2009; Grant et al., 2009; Beaty et al., 2010; Mangold et al., 2009; Wolf et al., 2015; Leslie et al., 2016a,b; Mostowska et al., 2018). However, the role of rare genetic variation in OFCs is still underway.

The Pittsburgh Orofacial Cleft Multiethnic GWAS (Leslie et al., 2016a,b) seeks to identify genetic variants that are associated with the risk of OFCs. This dataset includes a multi-ethnic cohort with 11,727 participants from 13 countries from North, Central or South America. Asia, Europe and Africa. Most of the participants were recruited as part of genetic and phenotyping studies coordinated by the University of Pittsburgh Center for Craniofacial and Dental Genetics and the University of Iowa. The study cohort includes OFC-affected probands with their family members, and controls without history of OFC. Affection status consists of cleft lip (CL) with or without palate (CL/P).

We performed standard data cleaning and quality control (see Leslie et al. (2016a)). We analyzed only Caucasian participants, and we kept rare variants with MAF in (0.001,0.05) and genotype call rate greater than 95%.

The final sample consisted of 1411 individuals, among which there were 835 controls, 309 cases without family history and 267 cases with family history. We did not include any covariates except for 5 principal components of ancestry (see Leslie et al. (2016a) for the details on the Principal Components Analysis). We applied ordinal GAMuT using linear kernel to measure pairwise phenotypic similarity. We also ran SKAT and burden tests, with the typical weights defined in Wu et al. (2011). For GAMuT, we used a weighted linear kernel (with the weighting scheme in Wu et al. (2011)) to measure pairwise genotypic similarity.

#### Data availability statement

The URLs for software: https://github.com/crsl4/ordinal-gamut and http://www.genetics.emory.edu/labs/epstein/software. The dataset title and accession number for dbGaP are “Center for Craniofacial and Dental Genetics (CCDG): Genetics of Orofacial Clefts and Related Phenotypes”, “dbGaP Study Accession: phs000774.v2.p1”.

## 3 Results

### 3.1 Type I Error Simulations

Figure 1 shows the quantile-quantile (QQ) plots of 10,000 null simulations with different subjects per group, target disease prevalence, and *λ* values. We show the comparison with ordinal GAMuT, SKAT and burden test. All methods compared properly control the type I error.

**Figure 1:**
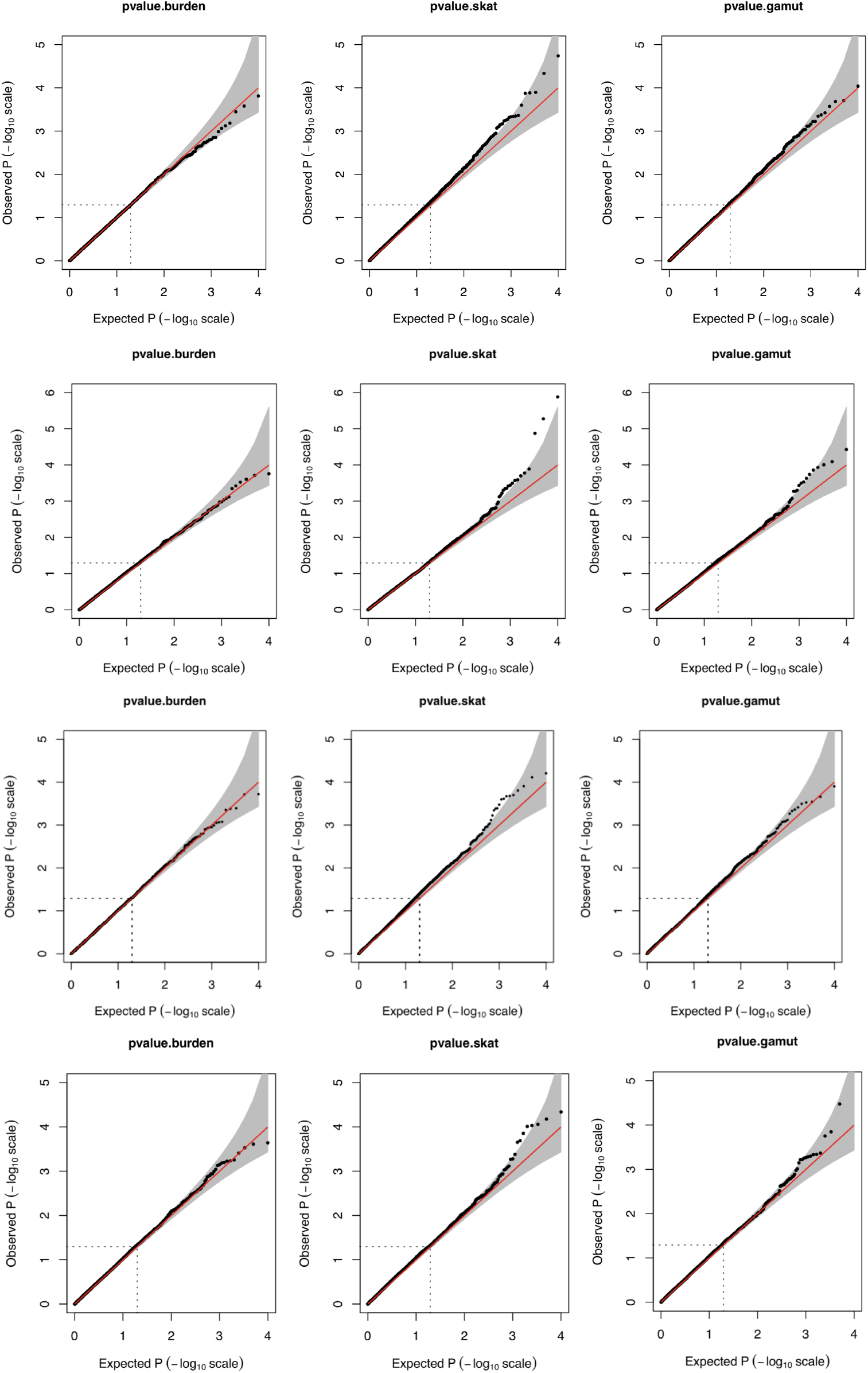
Q-Q plots of p-values for gene-based tests of rare variants for three methods: burden test, SKAT, and the ordinal GAMuT here proposed. Simulated datasets (10,000) assumed a 10kb region and rare variants defined as those with MAF <3%. **Top:** 750 subjects per group, disease prevalence of 0.01 and *λ* = 2. **Middle Top:** 750 subjects per group, disease prevalence of 0.05 and *λ* = 2. **Middle Bottom:** 750 subjects per group, disease prevalence of 0.01 and *λ* = 4. **Bottom:** 750 subjects per group, disease prevalence of 0.05 and *λ* = 4.

### 3.2 Power Simulations

Now, we compare the power of ordinal GAMuT with SKAT and burden test (Figure 2). The power was estimated by computing the proportion of p-values less than the significance level (*α* = 5 × 10^−5^ for effect sizes of 4 and 6) out of 1000 replicates per scenario and model. For these power simulations, we use different effect sizes in figures (*C* = 4,6). Columns refer to the conditional recurrence risk ratio *λ* = 2,4,8, and rows refer to disease prevalences 0.01,0.05. We compare the empirical power for sample sizes of *N* = 400,750,1000,1500 subjects per group. First, we note that we observe an increased number of causal variants in cases with family history compared to controls (see Supplementary Materials). Our method (ordinal GAMuT) outperformed the burden test (Li and Leal, 2008) and SKAT (Wu et al., 2011), with power increasing as sample size, recurrence risk, and effect size increased. Our method is more powerful given that other methods merge two clearly distinct groups: cases with and without family history, and thus, they cannot exploit the information present in the enrichment of causal variants in the cases with family history. Our ordinal approach models reality better by explicitly separating these two groups that have distinct genetic characteristics.

**Figure 2:**
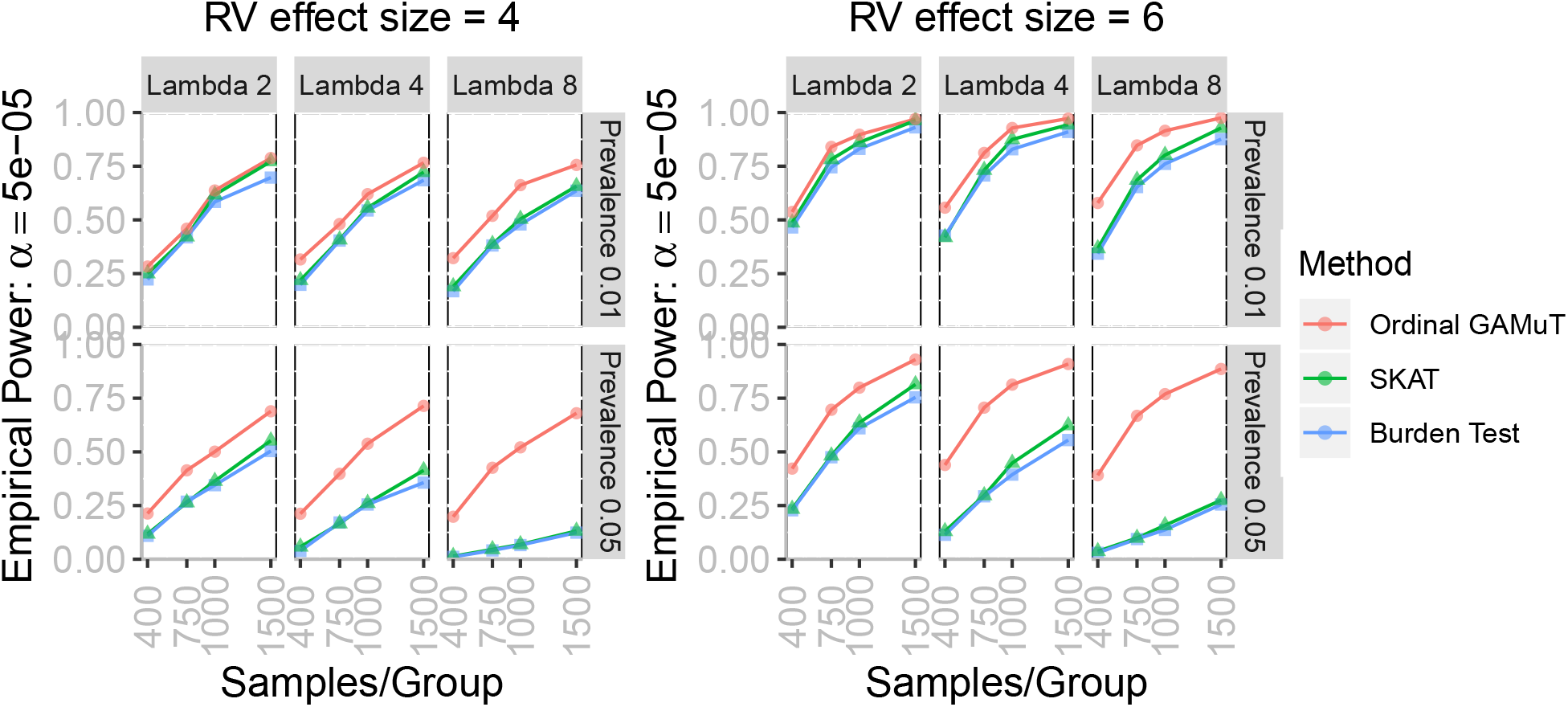
Power for gene-based testing comparing three methods: burden test (blue, square), SKAT (green, triangle) and ordinal GAMuT (red, circle). We compared two disease prevalences 0.01, 0.05 (rows), different conditional recurrence risk ratio *λ* = 2,4,8 (columns).

### 3.3 Analysis of Pittsburgh Orofacial Cleft Multiethnic GWAS

We applied our method to a Pittsburgh Orofacial Cleft (POFC) Multiethnic GWAS (Leslie et al., 2016a), (Leslie et al., 2016b) with 1,411 Caucasian subjects (267 cases with family history of clefting (up to third degree relatives), 309 cases without family history and 835 controls) and 61,671 variants used for annotation with Bystro (Kotlar et al., 2018). We filtered rare variants with MAF [0.001,0.05], and filtered genes to having minimum 4 rare variants, which resulted in 5,137 gene tests. We tested the association between the 5,137 genes and CL or CL/P status, adjusting for principal components for population structure. We compared our results (ordinal GAMuT) with the burden test and SKAT approach. Neither of the methods show any p-value inflation (Fig. 3).

**Figure 3:**
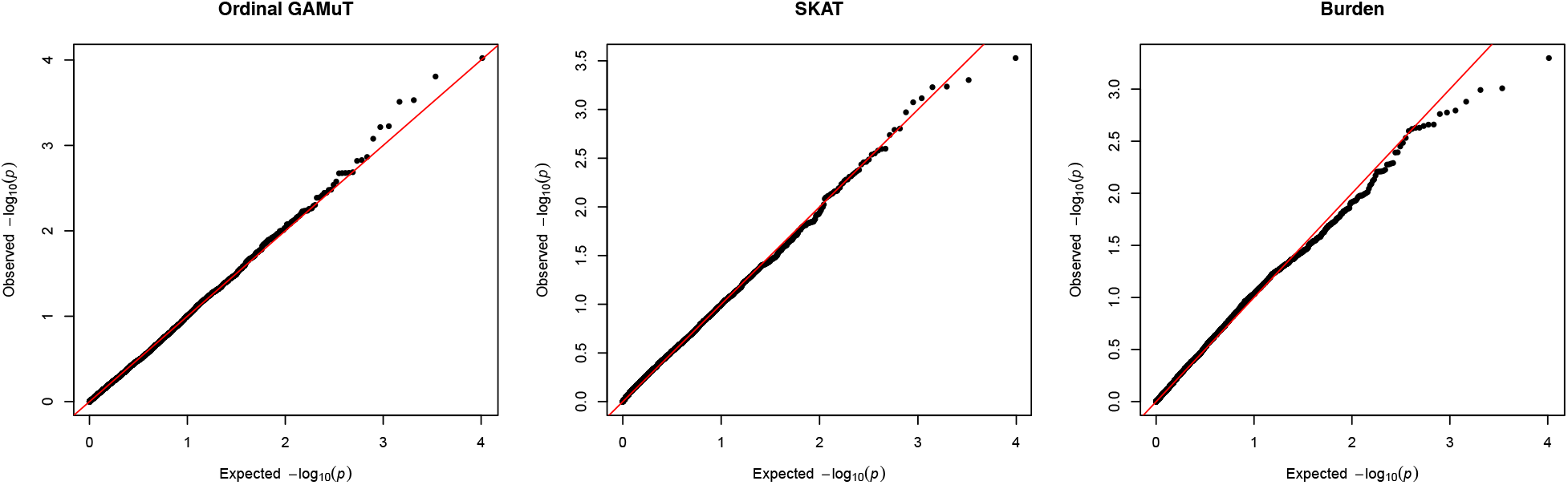
Q-Q plots of p-values for gene-based tests of rare variants for three methods: burden test (Li and Leal, 2008), SKAT (Wu et al., 2011) and the ordinal GAMuT here proposed in the GWAS of Pittsburgh Orofacial Cleft Multiethnic.

None of the methods identified any genes significantly associated with CL/P. However, ordinal GAMuT identified one gene (GRHL2) on chromosome 8 that passes the suggestive significance threshold (Fig. 4). GRHL2 is in the same gene family as GRHL3, which is a transcription factor that causes syndromic forms of clefting and is associated with nonsyndromic clefting in other GWAS (Leslie et al., 2016a,b; Carpinelli et al., 2017; Peyrard-Janvid et al., 2014).

**Figure 4:**
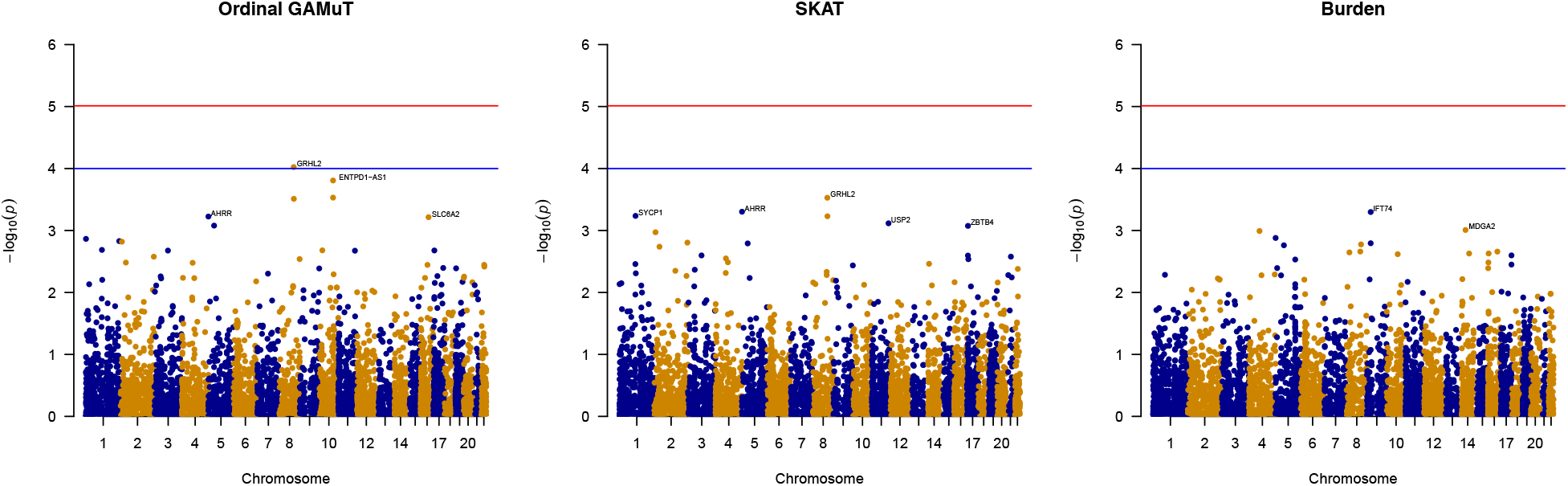
Gene-based test on Pittsburgh Orofacial Cleft (POFC) Multiethnic GWAS using burden, SKAT and ordinal GAMuT approach. Manhattan plots for each of the three tests. Red line: genome-wide significance level (− log_10_(0.05/5137) = 5.0117). Blue line: suggestive level (− log_10_(1 × 10^−4^) = 4).

## 4 Discussion

Standard GWAS methods for case-control studies usually define a disease outcome as a dichotomous phenotype. This phenotype ignores family history of disease, even if this information is available in the dataset at hand. Given that cases with a family history of disease can be enriched for risk variation relative to sporadic cases and may represent a source of case heterogeneity, incorporating family history is expected to increase power to detect genetic variants associated with disease.

We introduce an extension to the GAMuT method (Broadaway et al., 2016) to incorporate family information to enhance case-control association studies. This approach converts the usual binary phenotype of case-control status into an ordinal phenotype with three levels: cases with family history, cases without family history and controls, and it allows adjustment for covariates. Even though we do not include controls with family history, this ordinal approach can easily be extended to the case of four categories: controls with and without family history, and cases with and without family history by considering an ordinal phenotype with 4 levels. Finally, just as the standard GAMuT test, the ordinal GAMuT obtains analytic p-values from Davies’ method (Davies, 1980) which is computationally efficient, allowing the analysis of datasets in the genomic scale.

Simulation studies of rare variant sets showed that our ordinal GAMuT method is more powerful compared to usual gene-based tests like burden test (Li and Leal, 2008) and SKAT (Wu et al., 2011), possibly due to the fact that subjects with family history are more enriched for rare causal variants. Applying our method to Pittsburgh Orofacial Cleft Multiethnic GWAS (Leslie et al., 2016a,b), we identified a gene (GRHL2) (not previously reported) to suggestively associate with cleft lip and palate phenotypes. GRHL2 is in the same gene family as GRHL3, which is a transcription factor that causes in syndromic forms of clefting and was found to be associated with nonsyndromic clefting in GWAS (Leslie et al., 2016a,b; Carpinelli et al., 2017; Peyrard-Janvid et al., 2014). Burden and SKAT on these same phenotypes (figure 4) failed to identify any significant or suggestive genes. Among the weaknesses of the proposed method, extra care should be taken if there is a small cell count of cases with family history in the dataset, or in highly unbalanced dataset in which one of the categories is highly dominant in frequency compared to the other categories.

We envision two main future extensions of ordinal GAMuT: 1) to include information of more nuanced definitions of family history, and 2) to use disease liability as continuous phenotype instead of a categorical phenotype. Regarding disease liability, options for enhanced outcome variables could involve conditional means from liability-threshold models which have the potential to increase the power to detect genetic variants that are associated with disease risk. In fact, the popularity of proportional odds can be related to its connection to a linear regression model on a continuous latent response (e.g. the liability score). That is, the ordinal variable *Y* is obtained from a latent continuous variable *Z* by *Y* = *k* if *c*_*k*−1_ < *Z* ≤ *c*_*k*_. Thus, current ordinal GAMuT which utilizes proportional odds model has a natural extension into linear regression of the latent phenotype of disease liability. In the liability scale, family history can then be modeled as joint liability scores with a covariance matrix defined by the heritability of the disease.

Perhaps here or in model definition, we should note that the proportional odds/“parallel” assumption may be relaxed, highlighting the flexibility of the phenotype entering GAMuT (and by extension, the flexibility of ordinal GAMuT) Similarly, can replace the proportional odds model with something like an ordinal continuation ratio model (different logit formulations)

Finally, ordinal GAMuT is not restricted to rare genetic variants. Similar analysis could be performed for gene-based analysis of common variation.

## Acknowledgements

Data for the Orofacial Cleft Multiethnic GWAS comes from samples provided by Kaare Christensen (University of Southern Denmark), Frederic W.B. Deleyiannis (University of Colorado School of Medicine, Denver), Jacqueline T. Hecht (McGovern Medical School and School of Dentistry UT Health at Houston), George L. Wehby (University of Iowa), Seth M. Weinberg (University of Pittsburgh), Jeffrey C. Murray (University of Iowa) and Mary L. Marazita (University of Pittsburgh). This work was supported by NIH grants GM117946 [MP,DG], HG007508 [MP], R00-DE025060 [EJL], X01-HG007485 [MLM], R01-DE016148 [MLM, SMW], U01-DE024425 [MLM], R37-DE008559 [JCM, MLM], R21-DE016930 [MLM], R01-DE012472 [MLM], R01-DE011931 [JTH], R01-DE011948 [KC], U01-DD000295 [GLW]; NIH contract to the Johns Hopkins Center for Inherited Disease Research: HHSN268201200008I.

## Supplementary Material

### Enrichment of Causal Variants

In Figure 5, we show that, as expected, the average number of causal rare variants is greater for the cases with family history, followed by cases without family history, and lastly for controls. This simulated dataset comprises of 1000 controls, 1000 cases without family history, and 1000 cases with family history for three levels of conditional recurrence risk ratios (columns: *λ* = 2,4,8) and 2 siblings as family history. The effect size was set as *C* = 2.

**Figure 5:**
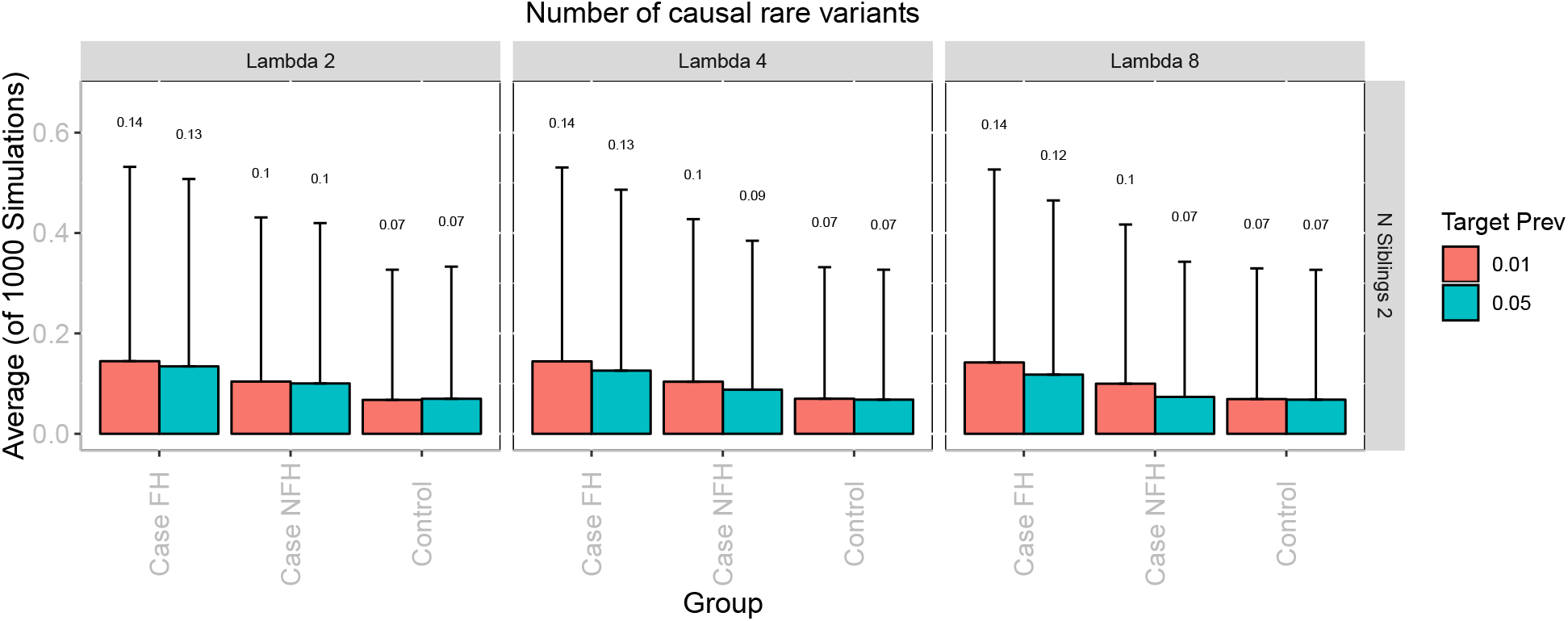
Average of 1000 simulations of number of causal rare variants (left) and probability of disease (right) in proband for three groups: controls, cases without family history, and cases with family history under two disease prevalences (red=0.01, blue=0.05), with one (top) or two (bottom) siblings, and three conditional recurrence risk ratios as columns.

## References

Beaty, T. H., Murray, J. C., Marazita, M. L., Munger, R. G., Ruczinski, I., Hetmanski, J. B., Liang, K. Y., Wu, T., Murray, T., Fallin, M. D., Redett, R. A., Raymond, G., Schwender, H., Jin, S.-C., Cooper, M. E., Dunnwald, M., Mansilla, M. A., Leslie, E., Bullard, S., Lidral, A. C., Moreno, L. M., Menezes, R., Vieira, A. R., Petrin, A., Wilcox, A. J., Lie, R. T., Jabs, E. W., Wu-Chou, Y. H., Chen, P. K., Wang, H., Ye, X., Huang, S., Yeow, V., Chong, S. S., Jee, S. H., Shi, B., Christensen, K., Melbye, M., Doheny, K. F., Pugh, E. W., Ling, H., Castilla, E. E., Czeizel, A. E., Ma, L., Field, L. L., Brody, L., Pangilinan, F., Mills, J. L., Molloy, A. M., Kirke, P. N., Scott, J. M., Arcos-Burgos, M., and Scott, A. F. (2010). A genome-wide association study of cleft lip with and without cleft palate identifies risk variants near mafb and abca4. Nature Genetics, 42:525 EP.

Birnbaum, S., Ludwig, K. U., Reutter, H., Herms, S., Steffens, M., Rubini, M., Baluardo, C., Ferrian, M., Almeida de Assis, N., Alblas, M. A., Barth, S., Freudenberg, J., Lauster, C., Schmidt, G., Scheer, M., Braumann, B., Bergé, S. J., Reich, R. H., Schiefke, F., Hemprich, A., Pötzsch, S., Steegers-Theunissen, R. P., Pötzsch, B., Moebus, S., Horsthemke, B., Kramer, F.-J., Wienker, T. F., Mossey, P. A., Propping, P., Cichon, S., Hoffmann, P., Knapp, M., Nöthen, M. M., and Mangold, E. (2009). Key susceptibility locus for nonsyndromic cleft lip with or without cleft palate on chromosome 8q24. Nature Genetics, 41:473 EP.

Broadaway, K. A., Cutler, D. J., Duncan, R., Moore, J. L., Ware, E. B., Jhun, M. A., Bielak, L. F., Zhao, W., Smith, J. A., Peyser, P. A., Kardia, S. L. R., Ghosh, D., and Epstein, M. P. (2016). A statistical approach for testing cross-phenotype effects of rare variants. American Journal of Human Genetics, 98(3):525–540.

Carpinelli, M., de Vries, M., Jane, S., and Dworkin, S. (2017). Grainyhead-like transcription factors in craniofacial development. Journal of Dental Research, 96(11):1200–1209. PMID: 28697314.

Davies, R. B. (1980). Algorithm as 155: The distribution of a linear combination of *χ*^2^ random variables. Journal of the Royal Statistical Society. Series C (Applied Statistics), 29(3):323–333.

De Rubeis, S., He, X., Goldberg, A. P., Poultney, C. S., Samocha, K., Cicek, A. E., Kou, Y., Liu, L., Fromer, M., Walker, S., Singh, T., Klei, L., Kosmicki, J., Shih-Chen, F., Aleksic, B., Biscaldi, M., Bolton, P. F., Brownfeld, J. M., Cai, J., Campbell, N. G., Carracedo, A., Chahrour, M. H., Chiocchetti, A. G., Coon, H., Crawford, E. L., Curran, S. R., Dawson, G., Duketis, E., Fernandez, B. A., Gallagher, L., Geller, E., Guter, S. J., Hill, R. S., Ionita-Laza, J., Jimenz Gonzalez, P., Kilpinen, H., Klauck, S. M., Kolevzon, A., Lee, I., Lei, I., Lei, J., Lehtimäki, T., Lin, C.-F., Ma’ayan, A., Marshall, C. R., McInnes, A. L., Neale, B., Owen, M. J., Ozaki, N., Parellada, M., Parr, J. R., Purcell, S., Puura, K., Rajagopalan, D., Rehnström, K., Reichenberg, A., Sabo, A., Sachse, M., Sanders, S. J., Schafer, C., Schulte-Rüther, M., Skuse, D., Stevens, C., Szatmari, P., Tammimies, K., Valladares, O., Voran, A., Li-San, W., Weiss, L. A., Willsey, A. J., Yu, T. W., Yuen, R. K. C., Study, D. D. D., for Autism, H. M. C., Consortium, U., Cook, E. H., Freitag, C. M., Gill, M., Hultman, C. M., Lehner, T., Palotie, A., Schellenberg, G. D., Sklar, P., State, M. W., Sutcliffe, J. S., Walsh, C. A., Scherer, S. W., Zwick, M. E., Barett, J. C., Cutler, D. J., Roeder, K., Devlin, B., Daly, M. J., and Buxbaum, J. D. (2014). Synaptic, transcriptional and chromatin genes disrupted in autism. Nature, 515(7526):209–215.

Epstein, M. P., Duncan, R., Ware, E. B., Jhun, M. A., Bielak, L. F., Zhao, W., Smith, J. A., Peyser, P. A., Kardia, S. L. R., and Satten, G. A. (2015). A statistical approach for rare-variant association testing in affected sibships. American Journal of Human Genetics, 96(4):543–554.

Grant, S. F. A., Wang, K., Zhang, H., Glaberson, W., Annaiah, K., Kim, C. E., Bradfield, J. P., Glessner, J. T., Thomas, K. A., Garris, M., Frackelton, E. C., Otieno, F. G., Chiavacci, R. M., Nah, H.-D., Kirschner, R. E., and Hakonarson, H. (2009). A genome-wide association study identifies a locus for nonsyndromic cleft lip with or without cleft palate on 8q24. The Journal of Pediatrics, 155(6):909–913.

Gretton, A., Fukumizu, K., Teo, C., Song, L., Schölkopf, B., and Smola, A. (2008). A kernel statistical test of independence. Adv. Neural Inf. Process. Systs, pages 585–592.

Hua, W.-Y. and Ghosh, D. (2015). Equivalence of kernel machine regression and kernel distance covariance for multidimensional phenotype association studies. Biometrics, 71(3):812–820.

Kosorok, M. R. (2009). On brownian distance covariance and high dimensional data. The annals of applied statistics, 3(4):1266–1269.

Kotlar, A. V., Trevino, C. E., Zwick, M. E., Cutler, D. J., and Wingo, T. S. (2018). Bystro: rapid online variant annotation and natural-language filtering at whole-genome scale. Genome Biology, 19(1):14.

Kwee, L., Liu, D., Lin, X., Ghosh, D., and Epstein, M. (2008). A powerful and flexible multilocus association test for quantitative traits. Am. J. Hum. Genet., 82:386–397.

Leslie, E. J., Carlson, J. C., Shaffer, J. R., Feingold, E., Wehby, G., Laurie, C. A., Jain, D., Laurie, C. C., Doheny, K. F., McHenry, T., Resick, J., Sanchez, C., Jacobs, J., Emanuele, B., Vieira, A. R., Neiswanger, K., Lidral, A. C., Valencia-Ramirez, L. C., Lopez-Palacio, A. M., Valencia, D. R., Arcos-Burgos, M., Czeizel, A. E., Field, L. L., Padilla, C. D., Cutiongco-de la Paz, E. M. C., Deleyiannis, F., Christensen, K., Munger, R. G., Lie, R. T., Wilcox, A., Romitti, P. A., Castilla, E. E., Mereb, J. C., Poletta, F. A., Orioli, I. M., Carvalho, F. M., Hecht, J. T., Blanton, S. H., Buxó, C. J., Butali, A., Mossey, P. A., Adeyemo, W. L., James, O., Braimah, R. O., Aregbesola, B. S., Eshete, M. A., Abate, F., Koruyucu, M., Seymen, F., Ma, L., de Salamanca, J. E., Weinberg, S. M., Moreno, L., Murray, J. C., and Marazita, M. L. (2016a). A multi-ethnic genome-wide association study identifies novel loci for non-syndromic cleft lip with or without cleft palate on 2p24.2, 17q23 and 19q13. Human molecular genetics, 25(13):2862–2872.

Leslie, E. J., Liu, H., Carlson, J. C., Shaffer, J. R., Feingold, E., Wehby, G., Laurie, C. A., Jain, D., Laurie, C. C., Doheny, K. F., McHenry, T., Resick, J., Sanchez, C., Jacobs, J., Emanuele, B., Vieira, A. R., Neiswanger, K., Standley, J., Czeizel, A. E., Deleyiannis, F., Christensen, K., Munger, R. G., Lie, R. T., Wilcox, A., Romitti, P. A., Field, L. L., Padilla, C. D., Cutiongco-de la Paz, E. M. C., Lidral, A. C., Valencia-Ramirez, L. C., Lopez-Palacio, A. M., Valencia, D. R., Arcos-Burgos, M., Castilla, E. E., Mereb, J. C., Poletta, F. A., Orioli, I. M., Carvalho, F. M., Hecht, J. T., Blanton, S. H., Buxó, C. J., Butali, A., Mossey, P. A., Adeyemo, W. L., James, O., Braimah, R. O., Aregbesola, B. S., Eshete, M. A., Deribew, M., Koruyucu, M., Seymen, F., Ma, L., de Salamanca, J. E., Weinberg, S. M., Moreno, L., Cornell, R. A., Murray, J. C., and Marazita, M. L. (2016b). A genome-wide association study of nonsyndromic cleft palate identifies an etiologic missense variant in grhl3. American journal of human genetics, 98(4):744–754.

Li, B. and Leal, S. M. (2008). Methods for detecting associations with rare variants for common diseases: application to analysis of sequence data. American journal of human genetics, 83(3):311–321.

Liu, J. Z., Erlich, Y., and Pickrell, J. K. (2017). Case–control association mapping by proxy using family history of disease. Nature Genetics, 49:325.

Mangold, E., Ludwig, K. U., Birnbaum, S., Baluardo, C., Ferrian, M., Herms, S., Reutter, H., de Assis, N. A., Chawa, T. A., Mattheisen, M., Steffens, M., Barth, S., Kluck, N., Paul, A., Becker, J., Lauster, C., Schmidt, G., Braumann, B., Scheer, M., Reich, R. H., Hemprich, A., Pötzsch, S., Blaumeiser, B., Moebus, S., Krawczak, M., Schreiber, S., Meitinger, T., Wichmann, H.-E., Steegers-Theunissen, R. P., Kramer, F.-J., Cichon, S., Propping, P., Wienker, T. F., Knapp, M., Rubini, M., Mossey, P. A., Hoffmann, P., and Nöthen, M. M. (2009). Genome-wide association study identifies two susceptibility loci for nonsyndromic cleft lip with or without cleft palate. Nature Genetics, 42:24 EP.

McCullagh, P. and Nelder, J. (1989). Generalized linear models. Chapman & Hall, 2nd edition.

Mossey, P. A., Little, J., Munger, R. G., Dixon, M. J., and Shaw, W. C. (2009). Cleft lip and palate. The Lancet, 374(9703):1773–1785.

Mostowska, A., Gaczkowska, A., Zukowski, K., Ludwig, K., Hozyasz, K., Wójcicki, P., Mangold, E., Böhmer, A., Heilmann-Heimbach, S., Knapp, M., Zadurska, M., Biedziak, B., Budner, M., Lasota, A., Daktera-Micker, A., and Jagodzinski, P. (2018). Common variants in dlg1 locus are associated with non-syndromic cleft lip with or without cleft palate. Clinical Genetics, 93(4):784–793.

Peyrard-Janvid, M., Leslie, E. J., Kousa, Y. A., Smith, T. L., Dunnwald, M., Magnusson, M., Lentz, B. A., Unneberg, P., Fransson, I., Koillinen, H. K., Rautio, J., Pegelow, M., Karsten, A., Basel-Vanagaite, L., Gordon, W., Andersen, B., Svensson, T., Murray, J. C., Cornell, R. A., Kere, J., and Schutte, B. C. (2014). Dominant mutations in grhl3 cause van der woude syndrome and disrupt oral periderm development. American journal of human genetics, 94(1):23–32.

Sanders, S. J., Neale, B. M., Huang, H., Werling, D. M., An, J.-Y., Dong, S., Abecasis, G., Arguello, P. A., Blangero, J., Boehnke, M., Daly, M. J., Eggan, K., Geschwind, D. H., Glahn, D. C., Goldstein, D. B., Gur, R. E., Handsaker, R. E., McCarroll, S. A., Ophoff, R. A., Palotie, A., Pato, C. N., Sabatti, C., State, M. W., Willsey, A. J., Hyman, S. E., Addington, A. M., Lehner, T., Freimer, N. B., and (WGSPD), W. G. S. f. P. D. (2017). Whole genome sequencing in psychiatric disorders: the WGSPD consortium. Nature Neuroscience, 20(12):1661–1668.

Schaffner, S. F., Foo, C., Gabriel, S., Reich, D., Daly, M. J., and Altshuler, D. (2005). Calibrating a coalescent simulation of human genome sequence variation. Genome research, 15(11):1576–1583.

Schaid, D. (2010). Genomic similarity and kernel methods ii: methods for genomic information. Hum. Hered., 70:132–140.

Székely, G. J., Rizzo, M. L., and Bakirov, N. K. (2007). Measuring and testing dependence by correlation of distances. The Annals of Statistics, 35(6):2769–2794.

Teng, J. and Risch, N. (1999). The relative power of family-based and case-control designs for linkage disequilibrium studies of complex human diseases. ii. individual genotyping. Genome Research, 9(3):234–241.

Tessier, P. (1976). Anatomical classification of facial, cranio-facial and latero-facial clefts. Journal of maxillofacial surgery, 4:69–92.

Wolf, Z. T., Brand, H. A., Shaffer, J. R., Leslie, E. J., Arzi, B., Willet, C. E., Cox, T. C., McHenry, T., Narayan, N., Feingold, E., Wang, X., Sliskovic, S., Karmi, N., Safra, N., Sanchez, C., Deleyiannis, F. W. B., Murray, J. C., Wade, C. M., Marazita, M. L., and Bannasch, D. L. (2015). Genome-wide association studies in dogs and humans identify adamts20 as a risk variant for cleft lip and palate. PLOS Genetics, 11(3):e1005059–.

Wu, M., Kraft, P., Epstein, M., Taylor, D., Chanock, S., Hunter, D., and X., L. (2010). Powerful snp-set analysis for case-control genome-wide association studies. Am. J. Hum. Genet., 86:929–942.

Wu, M. C., Lee, S., Cai, T., Li, Y., Boehnke, M., and Lin, X. (2011). Rare-variant association testing for sequencing data with the sequence kernel association test. American journal of human genetics, 89(1):82–93.

Yee, T. (2010). The vgam package for categorical data analysis. Journal of Statistical Software, Articles, 32(10):1–34.

Zhang, K., Peters, J., Janzing, D., and Schölkopf, B. (2012). Kernel-based conditional independence test and application in causal discovery. CoRR, abs/1202.3775.

Zöllner, S. (2012). Sampling strategies for rare variant tests in case-control studies. European journal of human genetics : EJHG, 20(10):1085–1091.

